# Effective Assessments of a Short-duration Poor Posture on Upper Limb Muscle Fatigue before Physical Exercise

**DOI:** 10.1101/2020.03.11.986653

**Authors:** Lei Lu, Mark Robinson, Ying Tan, Kusal Goonewardena, Xinliang Guo, Iven Mareels, Denny Oetomo

**Author notes:** Correspondence: Ying Tan.

## Abstract

A forward head and rounded shoulder posture is a poor posture that is widely seen in everyday life. It is known that sitting in such a poor posture with long hours will bring health issues such as muscle pain. However, it is not known whether sitting in this poor posture for a short period of time will affect human activities. This paper investigates the effects of a short-duration poor posture before some typical physical activities such as push-ups. The experiments are set up as follows. Fourteen male subjects are asked to do push-ups until fatigue with two surface electromyography (sEMG) at the upper limb. Two days later, they are asked to sit in this poor posture for 15 mins with 8 sEMG sensors located at given back muscles. Then they do the push-ups after the short-duration poor posture. The observations from the median frequency of sEMG signals at the upper limb indicate that the short-duration poor posture does affect the fatigue procedure of push-ups. A significant decreasing trend of the performance of push-ups is obtained after sitting in this poor posture. Such effects indicate that some parts of the back muscles indeed get fatigued with only 15 minutes sitting in this poor posture. By further investigating the time-frequency components of sEMG of back muscles, it is observed that the low and middle frequencies of sEMG signals from the infraspinatus muscle of the dominant side are demonstrated to be more prone to fatigue with the poor posture. Although this study focuses only on push-ups, similar experiments can be arranged for other physical exercises as well. This study provides new insights into the effect of a short-duration poor posture before physical activities. These insights can be used to guide athletes to pay attention to postures before physical activities to improve performance and reduce the risk of injury.

## 1 INTRODUCTION

Poor posture is regarded as prolonged deviations from the neutral spine, and it usually can be characterized by the forward head posture, rounded shoulders, and increased thoracic kyphosis (Singla and Veqar, 2017; Wong and Wong, 2008). Due to bad habits, poor posture is commonly observed in various scenarios in daily life. It is reported that poor posture presents emerging health risks (Jia and Nussbaum, 2018), especially, the prolonged poor posture is usually associated with the use of smartphones and other portable devices, which are reported having increasing musculoskeletal problems (Jung et al., 2016).

It is reported that prolonged poor posture may result in body discomfort and myofascial pain syndrome by placing stress and excessive tension on the lumbar vertebrae (Swann, 2009). For example, the forward head posture affected neck extensor muscle thickness (Goodarzi et al., 2018), and the spinal misalignment had bad influence on back muscle strength and shoulder range of motion (Imagama et al., 2014). Meanwhile, some other studies showed that poor posture had negative impact on the musculo-skeletal system, caused localized muscle fatigue, and may affect physical function and level of abilities (Ahmad and Kim, 2018).

In order to evaluate and measure the effect of poor postures, advanced sensor technologies have been developed. Many sensors with corresponding signal-processing techniques have been used to investigate the fatigue process during physical activities. Examples include the motion capture-based sensors that can capture the kinematic features of humans(Bailey et al., 2018), accelerators with dynamic movements and the related metrics (Beato et al., 2019), surface electromyography (sEMG) sensors to measure the muscle activities (Edouard et al., 2018), etc. Among them, the sEMG signals are shown to have excellent ability on revealing muscle activity at any time instant during various movements and postures Blanc and Dimanico (2010). The sEMG signal can provide information to characterize muscle fatigue by means of changes in signal indicators (Marshall et al., 2018; Toro et al., 2019), including the mean absolute value (MAV) (Toro et al., 2019), root mean square (RMS) (Girard et al., 2018), mean frequency (MNF) and median frequency (MDF) (Bowtell et al., 2014) and so on. As the sEMG signal is usually noisy, other advanced signal processing techniques, such as the wavelet transform, have been used to get better time-frequency resolutions (Chowdhury et al., 2013).

With analyzing sEMG signal collected from back muscles, it was shown that keeping the poor posture with long time had impacts on the development of muscle fatigue (Jia and Nussbaum, 2018). For example, with 1-hour typing task, the neck and shoulder pain were observed significantly increased with forward head and thoracic kyphotic posture (Kuo et al., 2019); with 40 mins poor posture sitting, the trunk flexion and metrics of localized muscle fatigue were significantly increased (Jia and Nussbaum, 2018).

However, it is not clear whether a short-duration of poor posture will have some negative impact on humans. It is known that the development of muscle fatigue is a time-varying procedure, it is challenging to detect muscle fatigue from the sEMG signal with a short-time duration of static poor posture, as changes are invisible due to small signal-to-noise ratio in sEMG signals. Although physiotherapists think that a short period of poor posture will also have some negative impact on people, there is a lack of systematic study to support this believe. As physical activities can somehow speed up the muscle fatigue process, this work tries to provide some evidence that sitting in poor posture indeed affect the performance of physical activities after it by carefully designed experiments. In this work, the push-up is selected to represent a common physical activities due to its simplicity.

The experiment consists of two steps. The first step measures the baseline of push-ups with two sEMG sensors to detect the fatigue of upper limb muscles. The second step, 15-minute of a poor sitting posture is followed by the same push-up activities. The duration of short-period poor posture is suggested by the physiotherapist. In the second step, each participant wears 8 sEMGs sensors on his back muscles. The location of each sEMG sensor is suggested and verified by a physiotherapist.

The hypothesis is that sitting in poor posture for short time will affect the performance of the following physical activities. If this hypothesis is true, this suggests that some parts of back muscles are getting fatigue during short period of poor posture, then some advanced signal processing techniques can be used to remove the noises and further investigate the measured sEMG signal.

The experimental results from 14 healthy male subjects support our hypothesis from the statistics of sEMG measurements. Our results indicate that poor posture, even a short-duration one, will have negative impact on human activities. Therefore, people should be always careful of avoiding poor postures.

## 2 MATERIALS AND METHODS

### 2.1 Participants

Fourteen healthy male college students were recruited for the experiment, the detailed information of the participants is shown in Table 1, including age, height, weight, and body mass index (BMI). All subjects were right-hand dominant. The participants were informed about the purpose and content of the experiment, and a written consent was obtained prior to the study. The project was approved by the Human Research Ethics Committee of the University of Melbourne (#1954575).

**Table 1.**
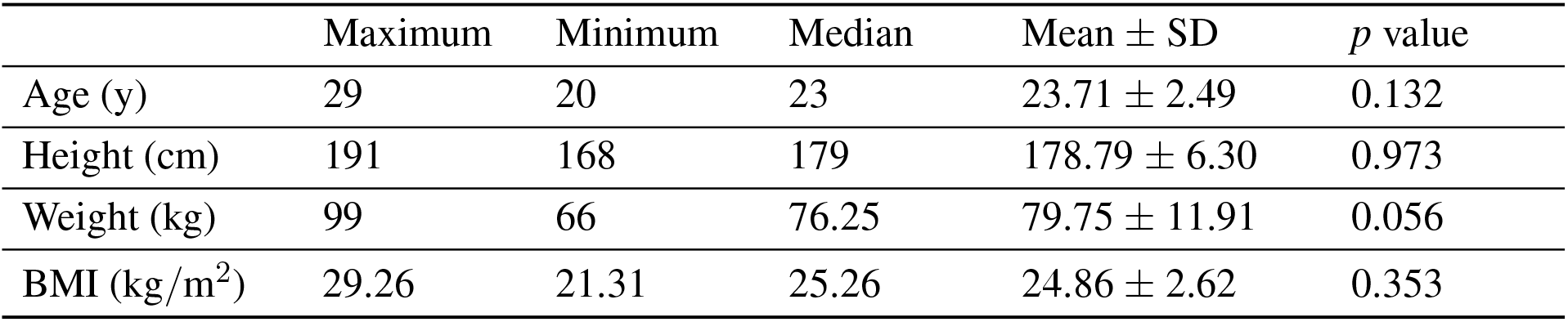
Characteristic information of the participants

### 2.2 Experimental Setup

The procedures of experimental setup are illustrated in Figure 1. As shown in Figure 1(A), the physical exercise of push-up is selected in our experiment as suggested by the physiotherapist. It is believed that similar performance can be observed if other physical activities are used.

**Figure 1.**
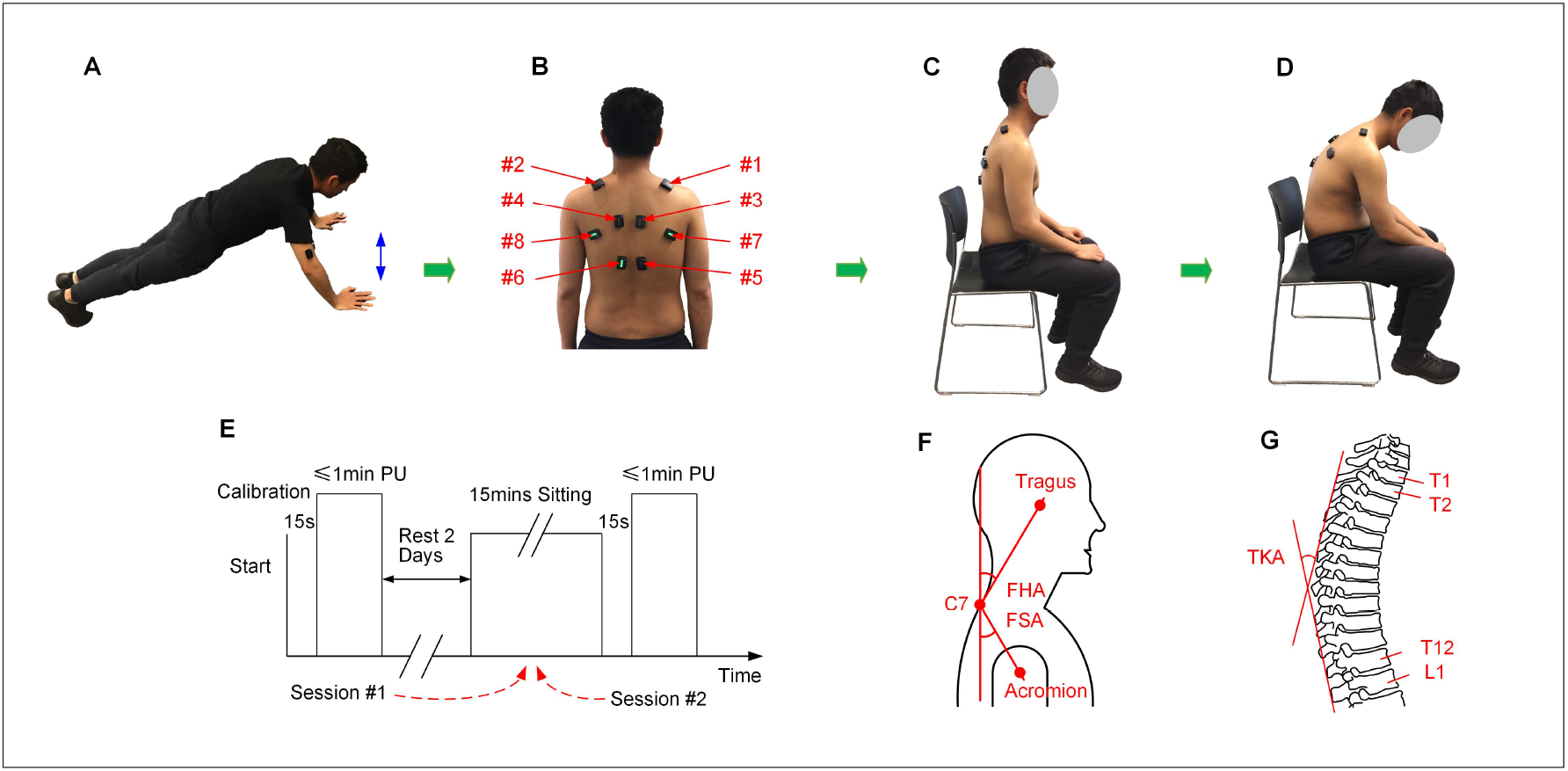
The experimental setup, (**A**) the physical exercise of push-up, (**B**) locations for the eight sEMG sensors, (**C**) illustration of the natural posture, (**D**) illustration of the poor posture, (**E**) the experimental procedure, (**F**) measurement of the FHA and FSA (Thigpen et al., 2010), (**G**) measurement of the TKA (Lewis and Valentine, 2010). *PU: Push-up*.

As discussed in Section 1, it is difficult to detect the small changes during poor posture sitting with a short time, the experiment consists of two sessions of push-ups (Figure 1(E)) to evaluate the influence of poor posture. During the first session, the participants were instructed to finish the exercise of push-up in their normal condition. In this session, the participants are asked to do push-ups for 1 minute or until the participants exhausted when it is less than 1 minute. The sEMG senors were calibrated before each experiment.

Before conducting the second experiment session, the participants were asked to have at least two days rest for muscle recovery. During the second session, the participants were first instructed to sit with the poor posture, which was illustrated in Figure 1(D): sitting with forward head, increased thoracic kyphosis, and rounded shoulder posture. The participants were advised to keep the poor posture for 15 minutes with minimal posture adjustments or in-chair movement. During the experiment, each subject was allowed to ’free his hands’, for example, the subject can play the mobile phone. After sitting with poor posture for 15 minutes, the participants were required to finish 1 minute push-up as described in the first session.

The DELSYS Trigno Biofeedback System (DELSYS Inc., Natick, MA, USA) was used to collect sEMG signals for analysis. Two sources of sEMG signals were used during the experiment, (1) the sEMG signals were collected from biceps brachii and triceps brachii during push-up exercises in both the two sessions. (2) During the poor posture sitting, sEMG signals were collected from eight upper back muscles, including the right upper fibers of trapezius (#1, RUFT), left upper fibers of trapezius (#2, LUFT), right middle fibers of trapezius (#3, RMFT), left middle fibers of trapezius (#4, LMFT), right lower fibers of trapezius (#5, RLFT), left lower fibers of trapezius (#6, LLFT), right infraspinatus (#7, RI), and left infraspinatus (#8, LI). Locations of the eight sensors was illustrated in Figure 1(B). The location of each sEMG sensor was adjusted for each participant according to the suggestion of the physiotherapist. Meanwhile, three angles of the natural sitting and poor sitting postures were measured, including the forward head angle (FHA), the forward shoulder angle (FSA), and the thoracic kyphosis angle (TKA) (Thigpen et al., 2010; Lewis and Valentine, 2010). Illustration of the three angles were demonstrated in Figures 1(F) and (G), and the measurements were shown in Table 2. The normality tests of the absolute difference of the angles were performed.

**Table 2.** Measurement of angles for the natural posture and the poor posture (Mean ± SD)

|  | Natural Posture | Poor Posture | Absolute Difference | *p* value |
| --- | --- | --- | --- | --- |
| FHA (°) | 39.81 ± 8.31 | 78.74 ± 5.07 | 38.93 ± 9.80 | 0.658 |
| FSA (°) | 28.96 ± 2.66 | 15.70 ± 2.95 | 13.26 ± 1.89 | 0.140 |
| TKA (°) | 36.90 ± 6.23 | 59.39 ± 1.51 | 22.49 ± 5.49 | 0.124 |

### 2.3 Statistical Analysis

Standard statistical analysis was performed on the collected data. Prior to the statistical analysis, data normality was checked with the Shapiro-Wilk test. Depending on whether the data distribution is normal or not, paired t-test or Wilcoxon signed rank test was performed to compare the frequency changes of push-up exercise in the two experiment sessions. One-way Analyses of Variance (ANOVA) with repeated measures or Kruskal-Wallis test was employed to measure frequency changes among different levels (the 1^st^ min, 5^th^ min, 10^th^ min, and 14^th^ min) during the poor posture sitting. The Tukey post hoc test was used to investigate the significant difference between different levels. The significance value was set at 0.05, and the data was presented as mean value ± standard deviation.

## 3 DATA ANALYSIS AND RESULTS

### 3.1 Poor Posture Affected Upper Limb Muscle Fatigue

The signal processing procedures for the sEMG signal collected from upper limb muscles during push-up are shown as Figure 2. The raw sEMG signal recorded at triceps brachii during the pushup is shown as Figure 2(A). The signal is sampled with 2148 Hz and some standard signal processing techniques were applied to pre-process this signal. These include outlier removal (three standard deviations from the mean value) and filtering (the 4^th^ order Butterworth band-pass filter with the frequency range of 10-500 Hz). The filtered signal is then used for envelope analysis and separating bursts of muscle activity during each push-up. The procedures are as following, first, the RMS is calculated to smooth the signal with time span of 100 ms; secondly, the envelope analysis is carried out to find the local minimal value of the RMS signal, and the valid burst separation points are obtained by comparing the identified local minimal points with a threshold, which can be roughly estimated by the time duration of burst, in the present study, the threshold value is usually set as 1s depending on each participant. The envelope analysis and identified bursts of push-up are shown as Figure 2(B), it can be seen from the figure that total 20 bursts of push-up are efficiently identified.

**Figure 2.**
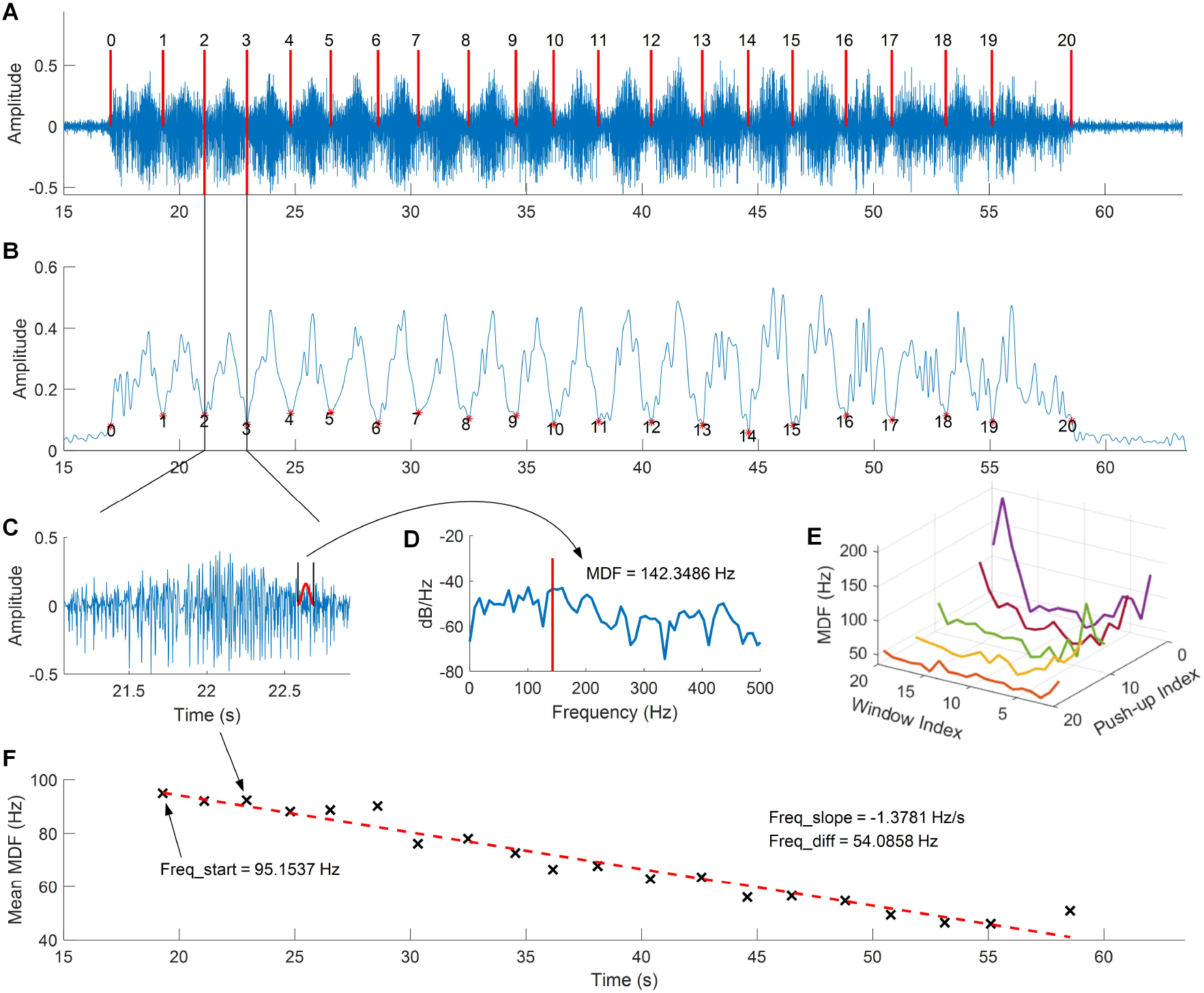
Procedures of the sEMG data analysis, (**A**) raw sEMG signal collected from the triceps brachii during push-up, (**B**) envelop analysis and push-up detection, (**C**) sEMG signal for a single push-up, (**D**) the spectral analysis for the moving hamming window segment, (**E**) median frequency curve of each push-up, (**F**) the mean median frequency and linear fitting of each push-up.

The MDF and MNF are then calculated for each burst, the Hamming window with length of 100 ms is used to obtain each segment in sEMG signal, the envelope of windowed signal is illustrated as Figure 2(C). Then the power spectral analysis for the segment is calculated as Figure 2(D), and the curve of MDF for each burst can be obtained with moving the Hamming window, which is illustrated as Figure 2(E) and the mean value of each curve is calculated for regression analysis. As shown in Figure 2(F), the linear regression is used to fit the calculated MDF values, and the frequency information is used for statistical analysis, including the slope of the linear curve, the difference between the the start and end frequencies, and the frequency difference percentage, which is calculated as the difference dividing by the start frequency.

The fatigue analysis with the mean MDF for pushup before and after the poor posture is demonstrated as Figure 3. It can be seen from Figure 3(A) that the slopes of the linear curves before and after poor posture are −2.506 and −1.3721, respectively. This result indicates the muscle fatigue has slower decreasing trend after the poor posture. Statistical analysis of the slope values with increasing the pushup numbers is shown in Figure 3(B), it can be seen from the figure that the slopes of the regression curves before poor posture are smaller than that after poor posture, while with push-up numbers of 10, the mean slopes of the regression curves before poor posture are significantly (*p* = 0.033) smaller than that after poor posture. The statistical analysis of the difference value between the start and the end frequency is shown as Figure 3(C), it can be seen from the figure that the mean value of the frequency difference before poor posture is larger than that after poor posture, with two significant values identified with 10 and 12 pushups. Figure 3(D) represents the statistical analysis for the percentage value of the frequency difference with a significant value obtained with 12 push-ups (*p* = 0.045), which indicates that the muscle fatigue process has a smaller decreasing percentage after poor posture.

**Figure 3.**
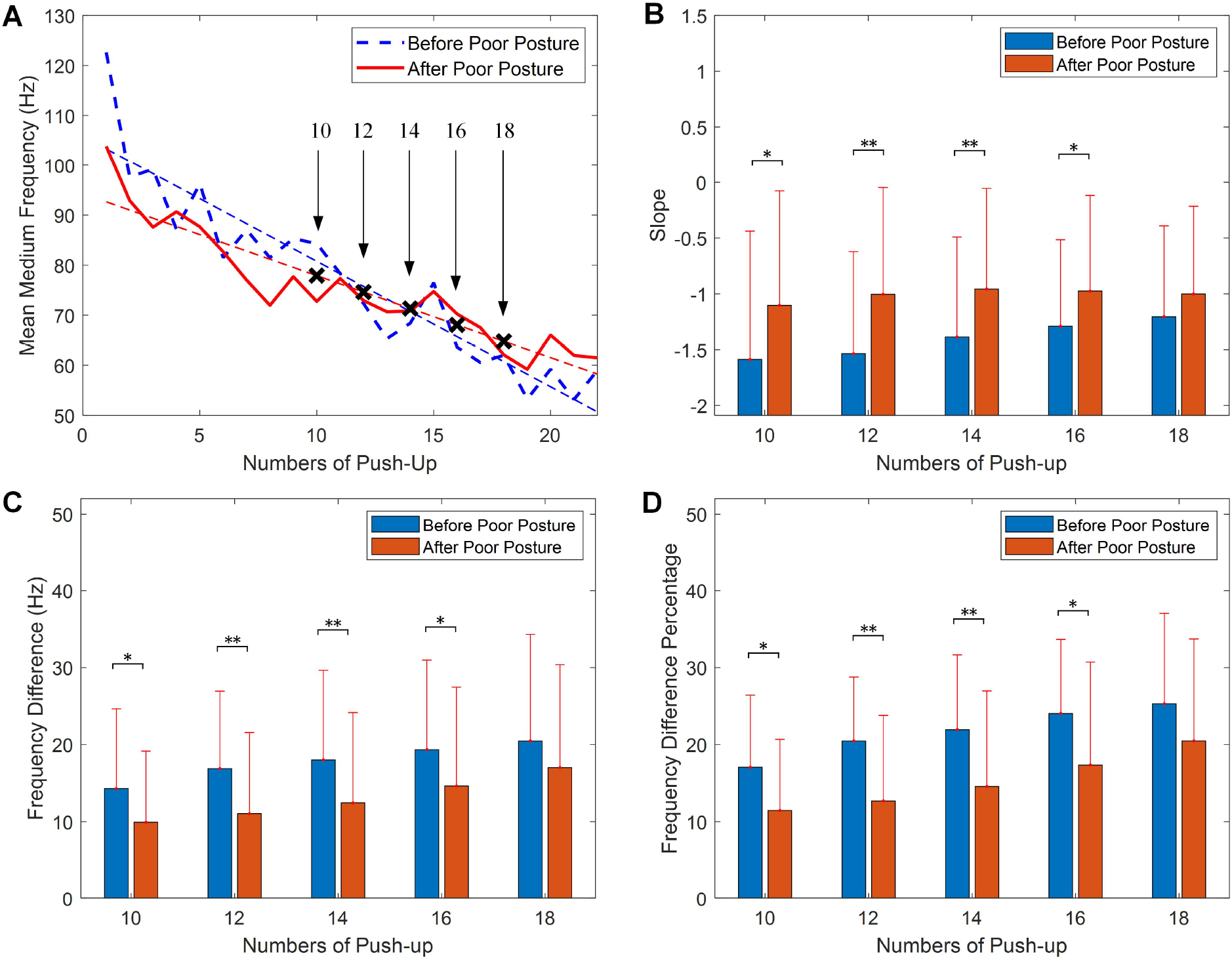
Fatigue analysis of push-up with mean median frequency (MDF), (**A**) comparison of linear regression before and after the poor posture, (**B**) statistical analysis of the slope values, (**C**) statistical analysis of the differences between the start frequency and the end frequency, (**D**) statistical analysis of the frequency difference percentage. (The data is presented as mean value ± standard deviation).

The similar results were obtained by calculating the mean MNF values of the triceps brachii. Figure 4 as indicated by the statistical analysis of frequency information of the mean MNF values. As shown in Figure 4(A) that the slope values of regression curve obtained for push-up before poor posture and after posture are −3.013 and −1.589. Figure 4(B) shows the comparison of mean slope values obtained for the two groups, the slope values obtained for 16 and 18 push-ups before poor posture are significantly (*p* = 0.033) smaller than that after poor posture. The similar comparison can be found in Figure 4(C). The comparison of statistical analysis for the percentage value of the frequency differences is shown as Figure 4(D), this figure shows that the mean percentage values of the frequency differences before poor posture are smaller than that of after posture.

**Figure 4.**
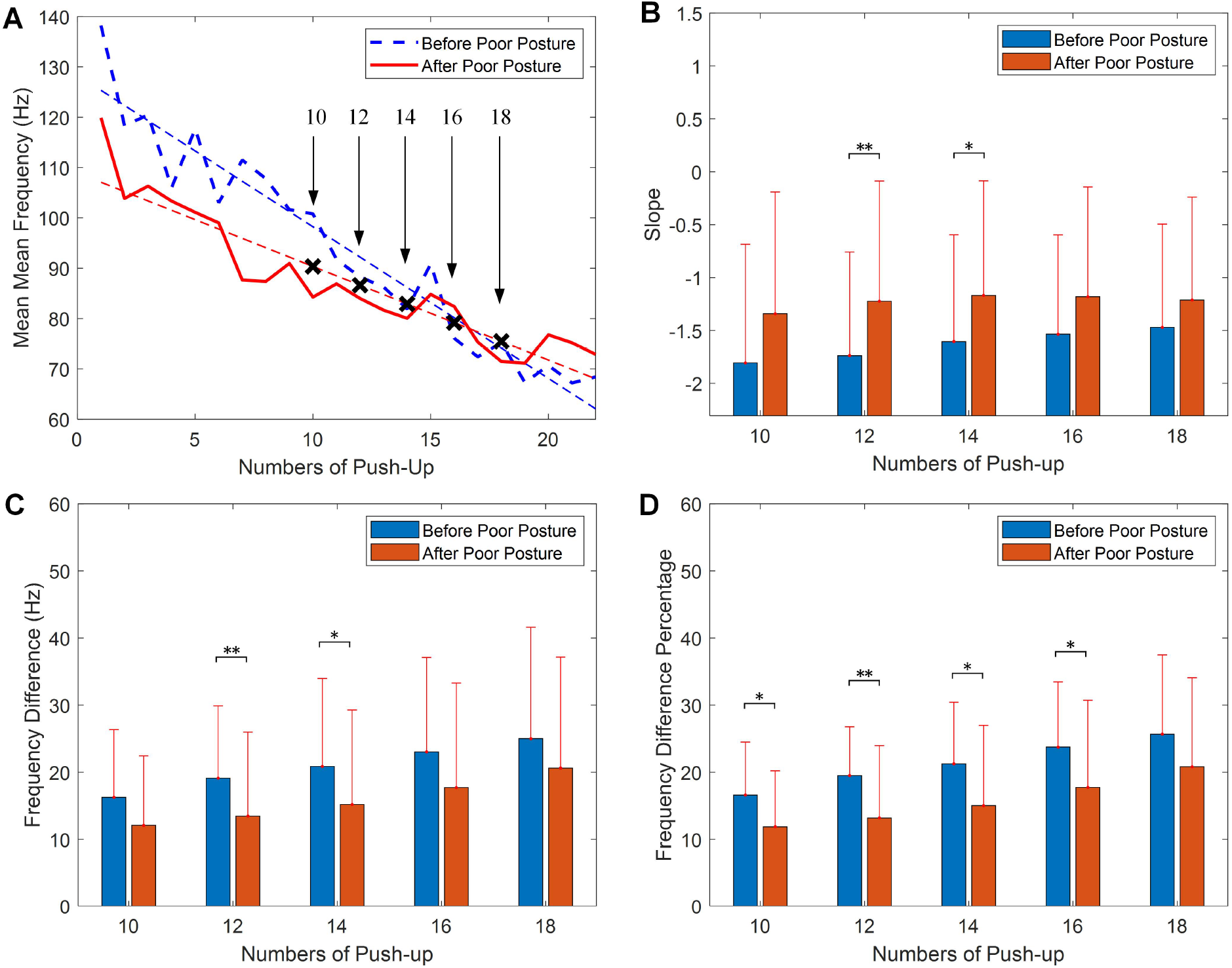
Fatigue analysis of push-up with mean mean frequency (MNF), (**A**) comparison of linear regression before and after the poor posture, (**B**) statistical analysis of the slope values, (**C**) statistical analysis of the differences between the start frequency and the end frequency, (**D**) statistical analysis of the frequency difference percentage. (Presented as mean value ± standard deviation).

### 3.2 Upper Back Muscle Fatigue Analysis

It can be seen from Section 3.1 that significant differences are observed for the push-up before and after the short-time poor posture sitting in terms of the frequency change in mean MDF and MNF, which indicates the poor posture has influence on the performance of physical exercise of the group. In this section, a detailed investigation of upper back muscles related to the poor posture is presented to evaluate the changes and sensitive frequencies of upper back muscles during the poor posture sitting.

As described in the experiment setup, eight wireless sensors are employed to collect sEMG signal from upper back muscles during the poor posture sitting. An illustration of the raw sEMG signal collected from the LLFT muscle during the 15 mins poor posture sitting is demonstrated in Figure 5(A). The widely used RMS and MDF features are calculated from the sEMG signal after band-pass signal filtering, which are time domain and frequency feature respectively. It can be seen from Figure 5(B) that there are heavy noise and fluctuation in the two features, and limit information can be obtained from the two features.

**Figure 5.**
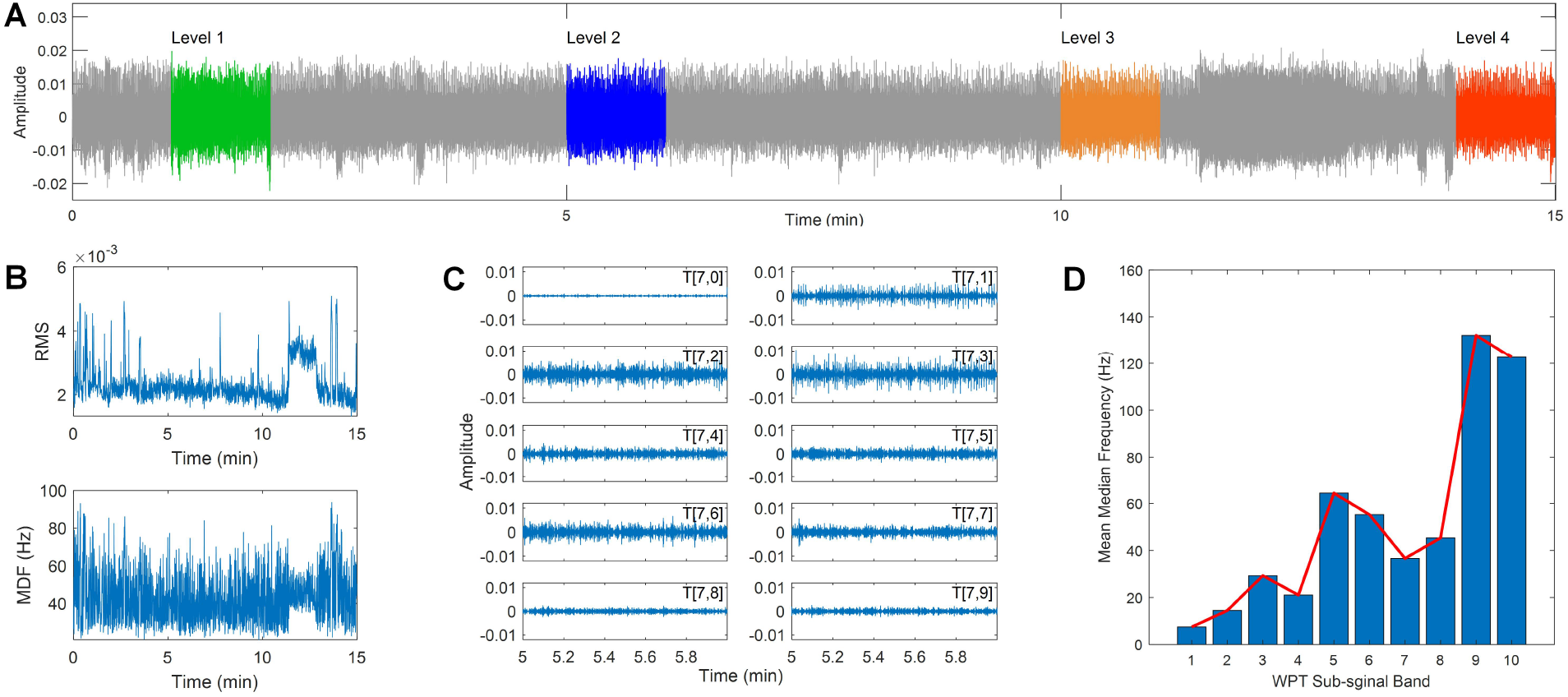
sEMG signal collected during static sitting and wavelet analysis, (**A**) the raw sEMG signal, (**B**) the RMS and median frequency features of the sEMG signal, (**C**) the sub-signal obtained from WPT decomposition, (**D**) the mean median frequency of each sub-signal.

As indicated in previous studies (Varrecchia et al., 2020; Chowdhury et al., 2013), the wavelet transform (WT) was a powerful technique on detecting small changes from noisy signal. The WT technique can obtain high resolution on analyzing sEMG signal by decomposing the raw signal into a series of sub-signals with different frequencies bands. Similar to the WT, the wavelet packet transform (WPT) is a generalization of WT and can obtain both low and high frequency information of the sEMG signal (Hekmatmanesh et al., 2019).

The WPT analysis involves two parameters, the wavelet function and the number of levels into which the signal will be decomposed. In the present study, the widely used wavelet families are employed for the sEMG signal analysis, including the Daubechies (db) wavelets, the Symlets (sym) wavelets, the Biorthogonal (bior) wavelets, and the Coiflets (coif) wavelets. The decomposition level was set as 7 to obtain a fine resolution in the frequency domain (Chowdhury et al., 2013; Kilby and Hosseini, 2004). The previous 10 subsignals of the original sEMG signal obtained by the WPT method is shown in Figure 5(C), and the corresponding median frequency of each sub-signal is demonstrated in Figure 5(D). It can be seen that the median frequency of the sub-signal indicates a gradually increasing trend with increasing the index number.

Four segments of the raw sEMG signal in the 1^st^ min, 5^th^ min, 10^th^ min, and 14^th^ min were used for the comparison study, which were denoted as level 1 to level 4. The WPT technique is used to extract sub-signals from the raw sEMG signal, and a moving window is applied to obtain the frequency spectrum for each sub-signal with the length of 1 s and the overlap rate of 0.5. The median frequency was calculated for each window frame and the mean value of median frequencies (MMDF) was calculated for all windows. Figure 6 demonstrates the calculated MMDF for all eight upper back muscles at the four time levels in different wavelet functions and sub-signals.

**Figure 6.**
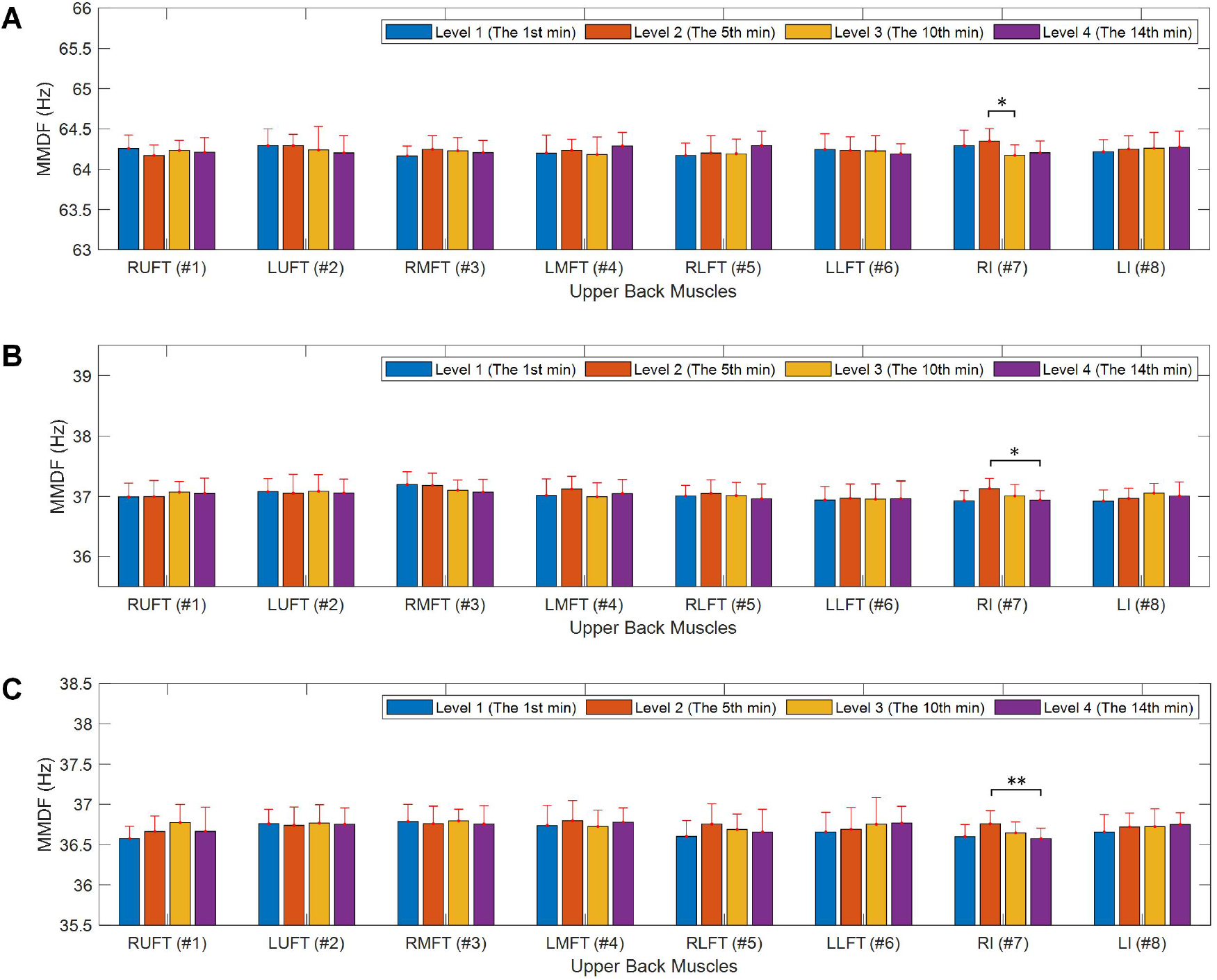
Upper back muscle fatigue analysis during the poor posture, (**A**) muscle fatigue analysis in the 5^th^ sub-signal with the coif4 wavelet, (**B**) muscle fatigue analysis in the 7^th^ sub-signal with the coif4 wavelet, (**C**) muscle fatigue analysis in the 7^th^ sub-signal with the coif2 wavelet.

It can be seen from Figure 6 that significant differences were observed for the RI muscle between four time levels in all the three sub-figures. Specifically, as demonstrated in Figure 6(A), with the coif4 wavelet the MMDF value of level 3 in the 5^th^ sub-signal and is observed smaller than the value of level 2 around 64 Hz significant value (*p* = 0.024). As demonstrated in Figure 6(B), the MMDF value of level 4 in the 7^th^ sub-signal is significantly smaller (*p* = 0.023) than the value of level 2 around 37 Hz. The similar frequency differences can be observed in Figure 6(C), the MMDF value of level 4 in the 7^th^ sub-signal obtained by the coif2 wavelet is smaller than the value of level 2 around 36.5 Hz with the significant value *p* = 0.010.

## 4 DISCUSSION

There are two major findings in this work, first, the short-time poor posture indeed has an effect on the performance of physical exercise. Compared with the performance of push-up before and after poor posture, the median frequencies of the two sessions have significant differences after 10 pushups in terms of the slope of the trend (*p* = 0.025), the frequency differences between the start and the end of the push-up (*p* = 0.025), and the rate of the frequency differences (*p* = 0.017). Second, It is confirmed from experiments and data analysis from sEMG sensors, some parts of back muscles are affected during the short time of poor posture with a clear indication of fatigue, which has not been observed in literature. As indicated in Figure 6, the median frequency of RI muscle is observed significant differences between different time levels.

Moreover, the obtained results using median frequency of sEMG signal collected from upper back muscles indicates that the wavelet transform especially the coif wavelet function has excellent performance on identifying the small frequency changes when signal-to-noise ration is small.

It is worthwhile to highlight that our analysis shows that sEMG signals of each muscle has it active frequency range. As shown in Figure 6, the median frequencies of the RI muscle around 36 Hz and 64 Hz are more sensitive. This finding is consistent with that in previous research (Chowdhury et al., 2013), which indicated that the sEMG signal induced by dynamic repetitive exertions are more sensitive to the lower frequencies.

It is noted that the short-duration poor posture is quite common in real life, for example. it might correspond to a sports break between two consecutive games. Another possible scenario is using one’s phone on the bus or train. Hence, the finding in this paper suggests that short-duration poor posture might lead to muscle fatigue of some parts of back muscles. After the poor posture, the performance of physical activities will be degraded (immediately and with latent effects). People should be mindful of their postures if they will undertake physical activities in the near future. The next step research will focus on the prevention of immediate and latent effects of muscle fatigue, which will provide know-how and awareness to prevent during sports and most importantly activities of daily living (ADLs).

## 5 CONCLUSIONS

This study contributed to the understanding of a particular short-duration poor sitting posture on the effect of the performance of physical exercise and the fatigue of back muscles. From statistics of 14 subjects, it was observed that the median frequencies of the sEMG signal of the triceps brachii has significant differences after poor sitting posture, and the median frequencies collected from the RI muscle in the upper back have been shifted in different time levels. The results suggest that the poor posture will lead to a quicker fatigue procedure of the following physical exercise as shown in the median frequency of the corresponding muscles. The results also demonstrated that the proposed frequency analysis techniques were able to identify the frequency range of the individual muscle, which can be used to show a clear shift of median frequencies of noisy sEMG signals in order to represent fatigue.

## CONFLICT OF INTEREST STATEMENT

Mr. Kusal Goonewardena is a sport physiotherapist for the Melbourne University Elite Athlete Unit, and CEO of the Elite Akademy Sports Medicine. All authors have no conflicts of interest related to this project.

## AUTHOR CONTRIBUTIONS

LL, XG, MR, KG and YT contributed the experiment design and data collection, LL and YT contributed the data analysis and drafting the manuscript, LL and MR performed the statistical analysis, KG, IM and DO contributed the manuscript revision and results interpretation. All the authors read and approved the manuscript.

## FUNDING

This work was supported in part by the National Natural Science Foundation of China (#51705106), funding from the Australian Wool Innovation Limited (AWI), and the International Postdoctoral Exchange Fellowship Program (#20170042).

